# ATM-Mediated Double-Strand Break Repair Is Required for Meiotic Genome Stability at High Temperature

**DOI:** 10.1101/2022.09.29.510164

**Authors:** Jiayi Zhao, Xin Gui, Ziming Ren, Huiqi Fu, Chao Yang, Qingpei Liu, Min Zhang, Wenyi Wang, Chong Wang, Arp Schnittger, Bing Liu

## Abstract

In eukaryotes, the conserved kinase Ataxia Telangiectasia Mutated (ATM) negatively regulates DNA double-strand break (DSB) formation and plays a central role in DSB repair. Here, by using cytogenetic approaches, we demonstrate that ATM also plays an essential role in protecting meiotic chromosome integrity in *Arabidopsis thaliana* at extreme high temperature. We determined the chromosome localization patterns of DSB formation proteins SPO11-1 and DFO during prophase I, both of which were disturbed by heat stress. Evaluation of the number of RAD51, DMC1, SPO11-1 and DFO protein foci in meiocytes of Arabidopsis *atm* mutant clarifies that ATM does not mediate the heat-induced reduction in DSB formation. Interestingly, meiotic spread analysis showed that chromosome fragmentation level was significantly increased in *atm* but was lowered in the *mre11* and *mre11 atm* mutants under high temperature, indicating that ATM-dependent meiotic chromosome integrity at high temperature relies on the functional MRE1-RAD50-NBS1 (MRN) complex. Moreover, contrary to the *rad51* and *mnd1* mutants, which exhibited enhanced meiotic chromosome integrity under heat stress, the *rad51 atm* and *mnd1 atm* mutants retained high levels of chromosome fragmentation at extreme high temperature. Furthermore, heat stress reduced chromosome fragmentation level in the *syn1* and *syn1 atm* mutants. Collectively, these data suggest that ATM-mediated DSB repair is required for meiotic genome stability in plants at extreme high temperature, which possibly acts in a RAD51-independent manner and relies on functional chromosome axis.

## Introduction

In eukaryotes, meiotic recombination generates novel combinations of genetic materials and creates genomic diversity; simultaneously, it safeguards the balanced segregation of homologs, which is crucial for the generation of viable gametes with halved ploidy (Wang and Copenhaver, 2018). Meiotic recombination is initiated by the formation of DNA double-strand breaks (DSBs) that are catalyzed by the DNA topoisomerase VIA subunit protein SPO11 together with other proteins within the DSB formation complex (in Arabidopsis, SPO11-1 and -2; PRD1, 2 and 3; DFO; MTOPVIB and PHS1 in Arabidopsis) (De Muyt et al., 2007; Grelon et al., 2001; Stacey et al., 2006; Vrielynck et al., 2021; Zhang et al., 2012). At the initiation of meiotic recombination, sister-chromatids are organized into a special structure called the chromosome axis, which plays important roles in regulating DSB formation and repair, homolog synapsis and recombination (Chambon et al., 2018; Cuacos et al., 2021; Ferdous et al., 2012; Lambing et al., 2022; Lambing et al., 2020). The RAD21-like cohesion protein REC8 (AtREC8/SYN1 in Arabidopsis) is a key chromosome axis factor that serves as a platform for the localization and function of other recombination proteins, with the *syn1* mutant having reduced DSB formation and impaired DSB repair and associated disrupted genome integrity (Bai et al., 1999; Cai et al., 2003; Lambing et al., 2020).

The MRE11-RAD50-NBS1 (MRN) complex processes the formed DSBs to generate 3’ overhang single-stranded DNAs (ssDNAs) and activates the downstream DNA damage response protein Ataxia Telangiectasia Mutated (ATM) to initiate DSB repair (Kurzbauer et al., 2012; Li et al., 2004; Sanchez-Moran et al., 2007; Syed and Tainer, 2018; Williams et al., 2010; Williams et al., 2007). ATM is a conserved kinase and regulates multiple meiotic recombination processes including DSB formation and repair, CO formation and distribution, and chromosome axis organization (Burma et al., 2001; Carballo et al., 2013; Garcia et al., 2003; Kurzbauer et al., 2021; Lange et al., 2011; Li and Yanowitz, 2019; Paiano et al., 2020; Plug et al., 1997; Yao et al., 2020). In different species, ATM negatively regulates DSB formation either by restricting the chromosome-localization of DSB formation proteins, or by engaging a chromosome axis-mediated DSB interference mechanism that affects DSB formation frequency near the generated DSB sites (Carballo et al., 2013; Cooper et al., 2014; Garcia et al., 2015; Kurzbauer et al., 2021; Lange et al., 2011; Paiano et al., 2020; Zhang et al., 2011).

ATM plays a central role in sensing and repairing DSBs (Shibata and Jeggo, 2021). A large-scale proteomic analysis in mammals revealed over 700 substrates for ATM and/or ATR in response to the DNA damage response, highlighting the complexity of the genetic network governed by ATM in regulating DSB repair (Matsuoka et al., 2007). However, the understanding of ATM substrates in regulating DSB repair in plants, especially during meiosis, is limited. In mammals, ATM activates and/or stabilizes downstream proteins such as RAD51 and XRCC3 to facilitate DSB repair (Ahlskog et al., 2016; Burma et al., 2001). RAD51 functions in DSB repair in mitotic and meiotic cells using sister-chromatids as templates; while DMC1 specifically acts in the homologous recombination (HR) pathway (Kurzbauer et al., 2012; Li et al., 2004; Sanchez-Moran et al., 2007). RAD51 serves as an accessory to DMC1 for catalyzing meiotic recombination-specific DSB repair by inhibiting the Structural Maintenance of Chromosomes 5/6 (SMC5/6) complex (Chen et al., 2021; Cloud et al., 2012; Kurzbauer et al., 2012). Loss of DMC1 function causes omitted crossovers (COs) formation, leading to univalent formation at diakinesis; while RAD51 dysfunction results in severe chromosome fragmentation due to impaired DSB repair (Cloud et al., 2012; Kurzbauer et al., 2012; Sanchez-Moran et al., 2007).

Global warming has now become one of the most serious climate factors that influence the living and evolution of land plants; and meiosis and HR in plants are prone to be affected by environmental temperature alterations (Bomblies et al., 2015; Liu et al., 2019; Modliszewski and Copenhaver, 2017). In Arabidopsis and in tree species, elevated temperatures have been found to induce meiotic restitution with resultant formation of unreduced gametes, interfere with CO formation and/or distribution, and affect spindle-dependent chromosome segregation which results in aneuploid gametes and sterility (De Storme and Geelen, 2020; Lei et al., 2020; Mai et al., 2019; Wang et al., 2017; Zhou et al., 2022). In *Arabidopsis thaliana*, a mildly high temperature within the fertility threshold (28°C) promotes the CO rate by enhancing the activity of the class I-type CO pathway (Lloyd et al., 2018; Modliszewski et al., 2018). However, higher temperatures at 32-34°C compromise CO formation (De Jaeger-Braet et al., 2022; De Storme and Geelen, 2020; Ning et al., 2021). When the temperature becomes extremely high that induces sterility (36-38°C), CO formation is completely abolished predominantly due to the interfered DSB formation and impaired homolog synapsis (Fu et al., 2021; Lei et al., 2020; Ning et al., 2021). However, the genetic mechanism that mediates the response of meiotic recombination in plants to elevated temperature has not yet been elucidated.

We have previously shown that the expression of ATM in Arabidopsis meiosis-staged flower buds is upregulated at high temperature (Ning et al., 2021), which suggests that the heat-induced reduction in DSB formation may be mediated through negative regulation by ATM. Here, by using a combination of cytological and genetic studies, we investigated the role of ATM in DSB formation and repair, homolog synapsis and CO formation in Arabidopsis at extremely high temperature. We showed that ATM does not mediate the heat-induced reduction in DSB formation; rather, it acts downstream of the MRN complex and probably regulates a RAD51-independent but SYN1-dependent DSB repair mechanism that protects meiotic chromosome integrity under heat stress. Overall, our study here demonstrates that ATM is required for maintenance of meiotic genome stability at extreme high temperature in *Arabidopsis thaliana*.

## Results

### Extreme high temperature reduces chromosome-localization of SPO11-1

In *Arabidopsis thaliana*, extremely high temperatures (36-38°C) interfere with the formation of double-strand breaks (DSBs) and suppress crossover (CO) formation (Fu et al., 2021; Ning et al., 2021). To address how heat stress reduces DSB formation, we examined the chromosome localization pattern of the DSB formation protein SPO11-1 in Arabidopsis plants under control (20°C) and extremely high temperature (37°C) conditions. A *pSPO11-1::SPO11-1-GFP* reporter was generated and introduced into the *spo11-1* mutant which has impaired DSB and CO formation (Hartung et al., 2007). The reporter fully rescued bivalent and tetrad formation as well as seed set in *spo11-1*, indicating that the recombinant SPO11-1-GFP was functional in catalyzing DSB formation and homologous recombination (HR) (Supplemental Fig. S1A-D). We analyzed the localization of SPO11-1-GFP on prophase I chromosomes by performing immunolocalization using an anti-GFP antibody (Fig. 1A). The chromosome axis protein SYN1, which is not sensitive to heat stress (Ning et al., 2021), was immunostained to indicate chromosome morphology. At 20°C, a low level of GFP foci accumulated in meiocytes at interphase, in which SYN1 displayed a punctate foci configuration (Fig. 1A). The number of GFP foci significantly increased beginning in leptotene, and peaked at zygotene and pachytene (Fig. 1A), and subsequently gradually decreased at diplotene (Fig. 1A). In line with a recent report, these findings revealed that in Arabidopsis, SPO11 increasingly accumulates until pachytene, when DSB number is significantly decreased and homologs have synapsed (Lambing et al., 2022).

**Figure 1.**
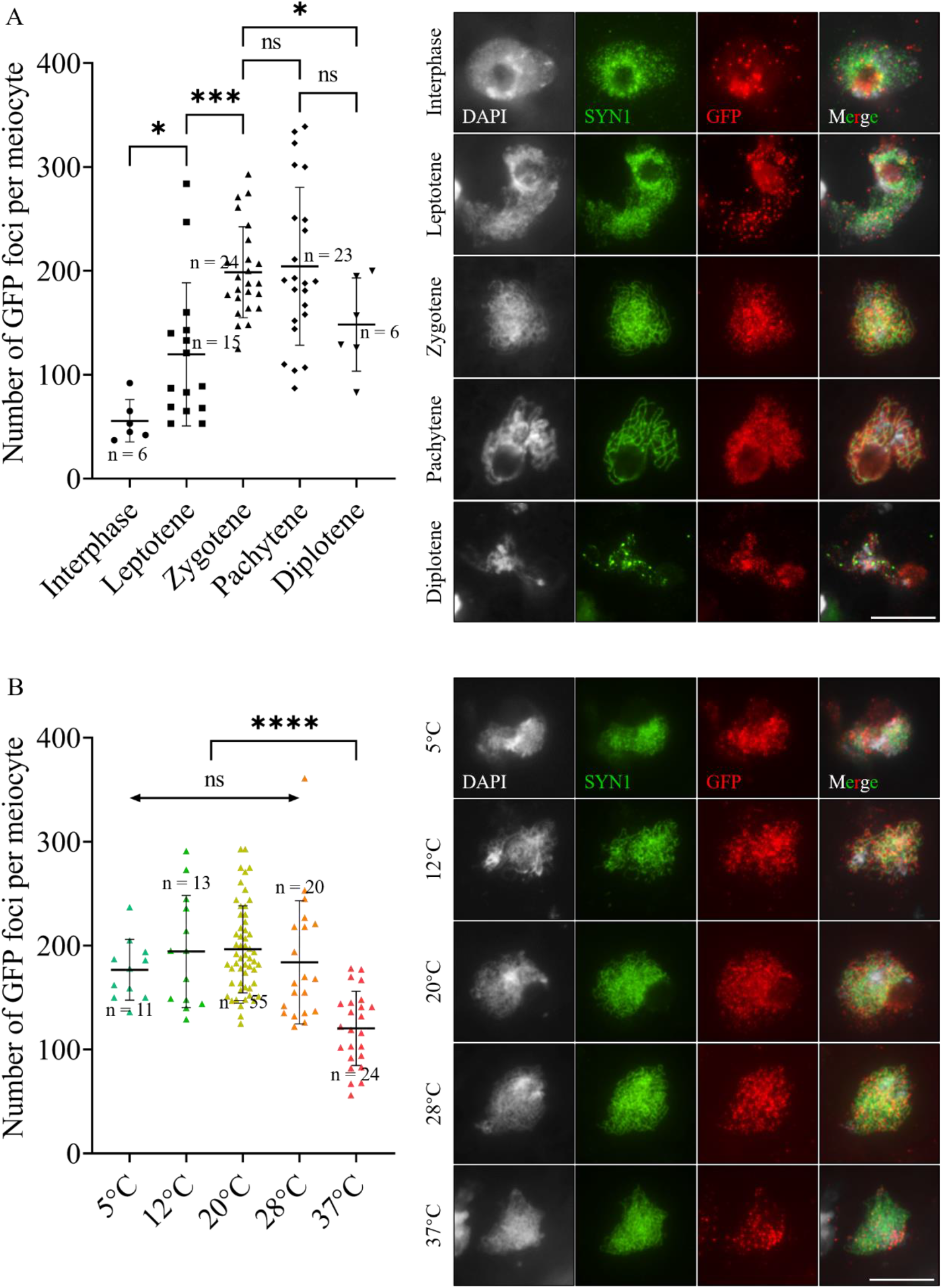
Analysis of the chromosome localization pattern of SPO11-1-GFP in *spo11-1-3* meiocytes expressing the *SPO11-1::SPO11-1-GFP* reporter (*spo11-1-3*^*SPO11-1::SPO11-1-GFP*^). A, Immunolocalization and quantification of SPO11-1-GFP in meiocytes at different prophase I stages in *spo11-1-3*^*SPO11-1::SPO11- 1-GFP*^ at 20°C. B, Immunolocalization and quantification of SPO11-1-GFP on zygotene chromosomes in *spo11-1-3*^*SPO11-1::SPO11-1-GFP*^ incubated at 20°C or shocked by 5, 12, 28 and 37°C for 24 h. Nonparametric and unpaired *t* tests were performed. n indicates the number of cells analyzed. **** indicates *P* < 0.0001;*** indicates *P* < 0.001; * indicates *P* < 0.05; ns indicates *P* > 0.05. Scale bars = 10 μm.

To elucidate the impact of increased temperature on chromosome localization of the SPO11 protein, we evaluated the number of SPO11-1-GFP foci on zygotene chromosomes in Arabidopsis plants incubated at 20°C or shocked by a series of elevated temperatures from 5, 12, 28 to 37°C for 24 h (Fig. 1B). We found that the number of GFP foci remained at a relatively stable level from 5°C, at which meiotic restitution can occur (De Storme et al., 2012; Liu et al., 2018), to 28°C, at which class I-type crossover (CO) rate increases (Lloyd et al., 2018; Modliszewski et al., 2018). In comparison, the number of GFP foci was significantly lowered at 37°C (Fig. 1B). According to the number of SPO11-1 protein foci, immunolocalization of the DSB repair protein DMC1 on zygotene chromosomes in Col-0 plants at 5, 12 or 28°C for 24 h showed numbers of DMC1 foci similar to those in plants incubated at 20°C, but the DMC1 foci number in plants at 37°C was significantly lowered (Supplemental Fig. S2). These data suggested that extremely high temperature interferes with the accumulation of SPO11-1 protein on prophase I chromosomes, which may induce a reduction in DSB formation.

### Localization pattern of DFO on prophase I chromosomes at increased temperature

To confirm that heat stress reduces DSB formation by interfering with chromosome localization of DSB formation proteins, we analyzed the chromosome localization pattern of DFO protein, which is also required for DSB formation in Arabidopsis (Zhang et al., 2012). Immunolocalization of DFO protein was performed using an anti-DFO antibody. Dual-immunolocalization of DFO and SYN1 in Col-0 grown at 20°C showed a relatively small number of punctate DFO foci in interphase meiocytes, which were localized on thin-thread chromosomes (Fig. 2A; Supplemental Fig. S3A). Subsequently, DFO foci started to accumulate at leptotene, retained at a relatively stable level until late pachytene, and then decreased at diplotene (Fig. 2A; Supplemental Fig. S3B-F). The observed colocalization of DFO foci and SYN1 signals (Supplemental Fig. S3) supported a direct interaction between DFO and chromosome axis proteins (Vrielynck et al., 2021). The *dfo* mutant, which has impaired homolog synapsis and bivalent formation (Supplemental Fig. S4) (Zhang et al., 2012), did not show any specific DFO foci (Supplemental Fig. S5), confirming that the antibody was reliable. After heat stress treatment, we observed a significantly reduced number of DFO foci on leptotene, zygotene and pachytene chromosomes (Fig. 2B), indicating that heat stress compromises the localization of DFO protein on prophase I chromosomes. Reverse transcription quantitative PCR (RT-qPCR) showed that the expression of *DFO* in meiosis-staged flower buds of Col-0 was reduced at extreme high temperature (Supplemental Fig. S6C), which may partially contribute to the observed reduction in DFO abundance in the meiocytes.

**Figure 2.**
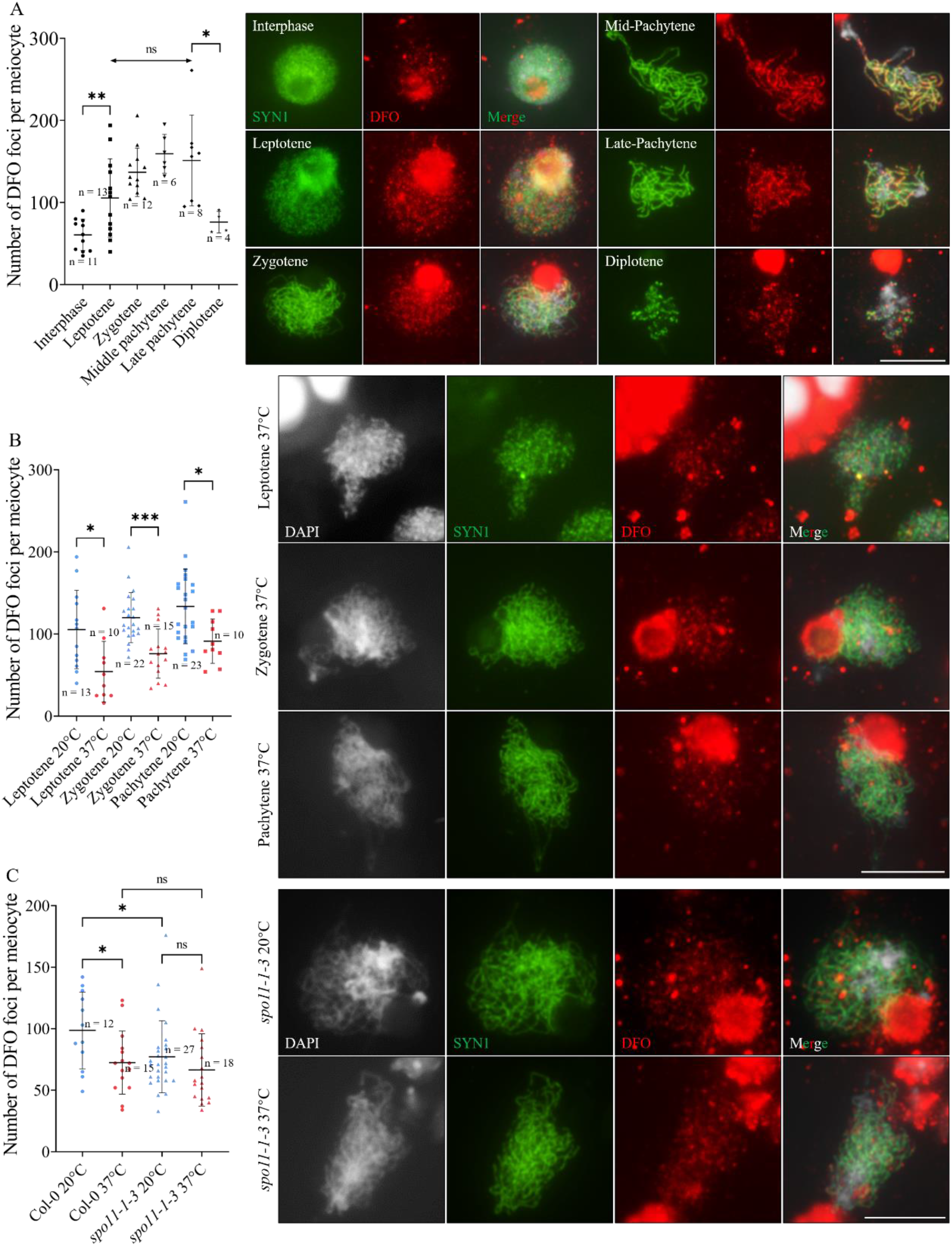
Analysis of chromosome localization of DFO in Col-0 and *spo11-1-3* at increased temperature. A, Immunolocalization and quantification of DFO in Col-0 meiocytes at different prophase I stages at 20°C. B, Immunolocalization and quantification of DFO on leptotene, zygotene and pachytene chromosomes in Col-0 at 37°C. C, Immunolocalization and quantification of DFO on zygotene chromosomes in Col-0 and *spo11-1-3* at 37°C. Nonparametric and unpaired *t* tests were performed. n indicates the number of cells analyzed. *** indicates *P* < 0.001; ** indicates *P* < 0.01; * indicates *P* <0.05; ns indicates *P* > 0.05. Scale bars = 10 μm.

In Arabidopsis, it has been shown that the chromosome-localization of SPO11-1 does not rely on the DFO protein (Vrielynck et al., 2021). To determine the epistatic relationship between the chromosome localization of SPO11-1 and DFO, we analyzed DFO foci formation in *spo11-1*. At 20°C, *spo11-1* showed a smaller number of DFO foci on zygotene chromosomes than Col-0 (Fig. 2C), suggesting that the normal chromosome localization of DFO is SPO11-1 dependent. Furthermore, we found that heat stress did not further reduce the number of DFO foci in *spo11-1* (Fig. 2C), which implied that heat stress may influence DFO localization dependently on SPO11-1.

### Heat stress reduces RAD51 and DMC1 foci in *atm*

In many organisms including Arabidopsis, the kinase Ataxia Telangiectasia Mutated (ATM) negatively regulates DSB formation through several mechanisms, including restricting the accumulation of DSB formation proteins (Carballo et al., 2013; Garcia et al., 2015; Kurzbauer et al., 2021; Paiano et al., 2020). The reduced chromosome localization of SPO11-1 and DFO proteins (Fig. 1B and Fig. 2C), and the increased expression of *ATM* at high temperature revealed by RT-qPCR (Supplemental Fig. S6B) (Ning et al., 2021) led us to hypothesize that the reduced DSB formation at high temperature is mediated by ATM. To clarify this hypothesis, we evaluated the DSB levels in Col-0 and *atm* under control and high temperature conditions. ɤH2A.X is the phosphorylated form of histone H2A.X that specifically localizes near DSB sites and thus is often used as a marker of DSBs (Burma et al., 2001). In Col-0, the number of ɤH2A.X foci can be used as an indicator of DSB levels, and the effect of heat stress on ɤH2A.X foci formation was found to be SPO11-1-dependent (Supplemental Fig. S7). However, since H2A.X phosphorylation is catalyzed by ATM, and Arabidopsis plants with impaired ATM function have significantly compromised ɤH2A.X foci formation (Supplemental Fig. S8) (Yao et al., 2020), we did not score the number of ɤH2A.X foci as the indicator of DSB number in *atm*. Instead, we performed immunolocalization of the DSB repair recombinases RAD51 and DMC1, which are also widely used as indirect DSB markers (Dereli et al., 2021; He et al., 2017; Kurzbauer et al., 2021). At 20°C, in line with other reports, there were more RAD51 and DMC1 foci in *atm* meiocytes than those in Col-0, indicating a higher number of DSBs or an interfered progressing of DSB repair when ATM was absent (Fig. 3A and B) (De Jaeger-Braet et al., 2022; Kurzbauer et al., 2021; Yao et al., 2020). After heat stress treatment, the numbers of both RAD51 and DMC1 foci were reduced in Col-0 and *atm* (Fig. 3A and B), suggesting that heat stress reduced DSB formation in *atm*.

**Figure 3.**
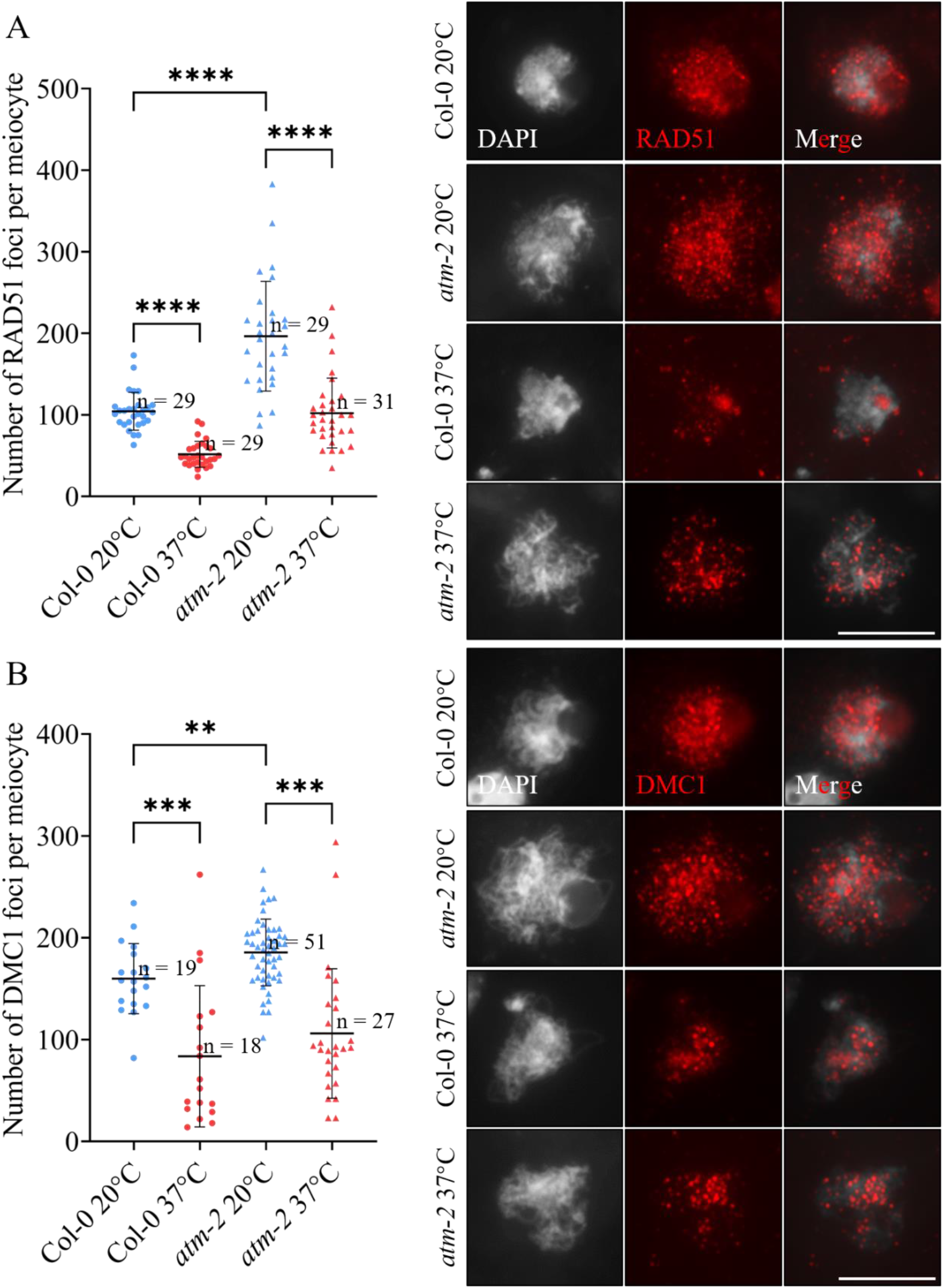
Quantification of RAD51 and DMC1 foci on zygotene chromosomes in Col-0 and *atm-2* at 20 and 37°C. A, Immunolocalization and quantification of RAD51 on zygotene chromosomes of Col-0 and *atm-2* at 20 and 37°C. B, Immunolocalization and quantification of DMC1 on zygotene chromosomes of Col-0 and *atm-2* at 20 and 37°C. Nonparametric and unpaired *t* tests were performed. n indicates the number of cells analyzed. **** indicates *P* < 0.0001; *** indicates *P* < 0.001; ** indicates *P* < 0.01. Scale bars = 10 μm.

### Heat stress reduces the number of SPO11-1 and DFO foci in *atm*

To confirm that heat stress reduces DSB formation in *atm*, we compared the effect of extreme high temperature on chromosome localization of the SPO11-1 and DFO proteins on zygotene chromosomes in Col-0 and *atm*. At 20°C, a higher number of SPO11-1-GFP and DFO foci was observed in *atm* than in Col-0 (Fig. 4A and B), suggesting a negative impact of ATM on the chromosome localization of these two DSB formation factors in Arabidopsis. At 37°C, the numbers of SPO11-1-GFP and DFO foci were lowered in both Col-0 and *atm* (Fig. 4A and B). These findings demonstrated that ATM does not mediate the heat-induced reduction in DSB.

**Figure 4.**
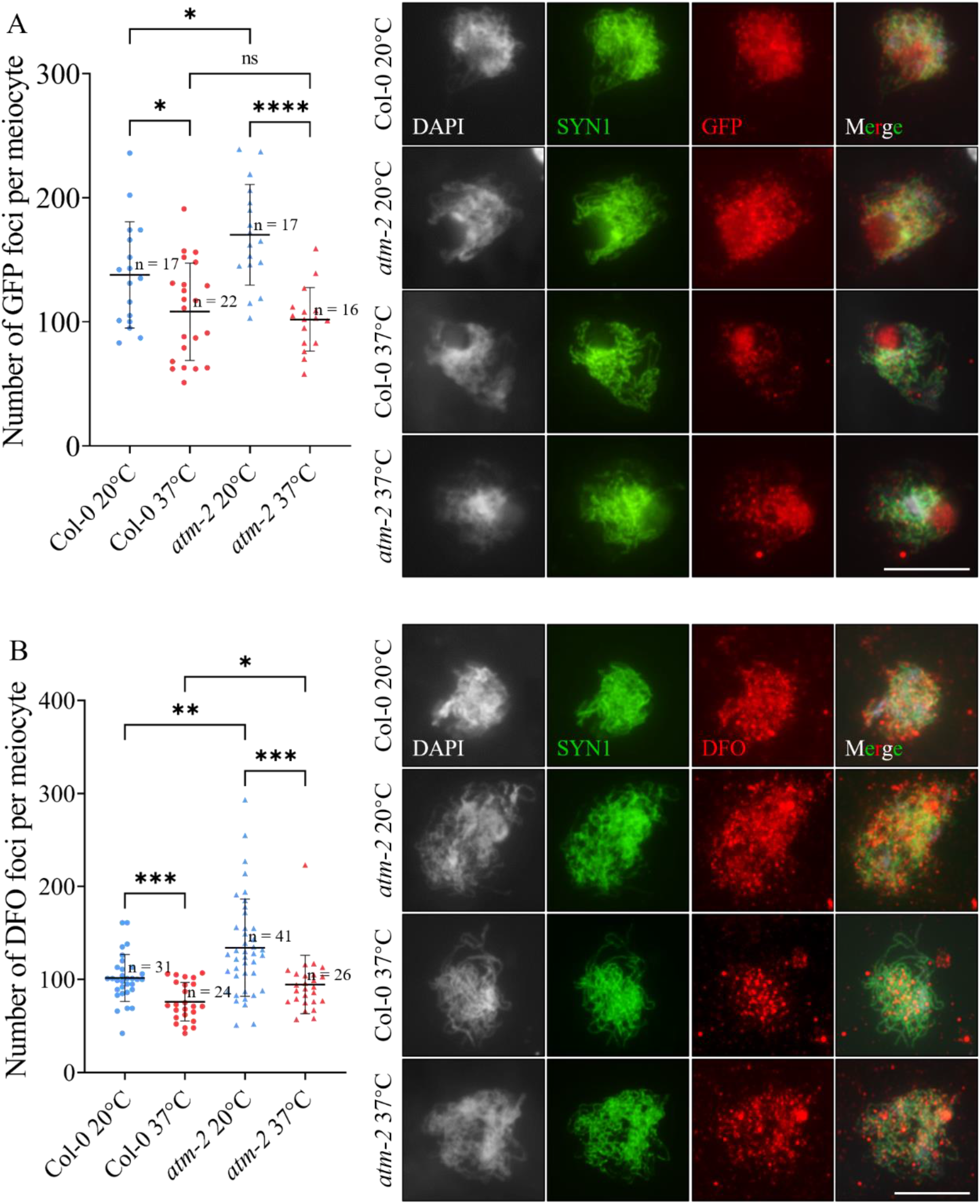
Analysis of chromosome-localization of SPO11-1-GFP and DFO in Col-0 and *atm-2* at 20 and 37°C. A, Immunolocalization and quantification of SPO11-1-GFP on zygotene chromosomes of Col-0 and *atm-2* at 20 and 37°C. B, Immunolocalization and quantification of DFO on zygotene chromosomes of Col-0 and *atm-2* at 20 and 37°C. Nonparametric and unpaired *t* tests were performed. n indicates the number of cells analyzed. **** indicates *P* < 0.0001; *** indicates *P* < 0.001; ** indicates *P* < 0.01; * indicates *P* < 0.05; ns indicates *P* > 0.05. Scale bars = 10 μm.

### Heat stress increases chromosome fragments in *atm* meiocytes

Dysfunction of ATM in Arabidopsis causes chromosome fragmentation due to interfered DSB repair (Kurzbauer et al., 2021; Yao et al., 2020). We thereafter questioned whether ATM also plays a role in DSB repair under heat stress. To this end, we performed meiotic spread analysis and evaluated the chromosome fragmentation level in *atm* meiocytes at metaphase I, anaphase I and metaphase II at increased temperature. In Col-0, heat stress induced the formation of univalents with disordered homolog segregation, while the chromosomes maintained complete integrity (Fig. 5A, metaphase I; B, anaphase I; C, metaphase II). Similarly, the *spo11-1* mutant did not show any chromosome fragmentation at either temperature (Fig. 5A, metaphase I; B, anaphase I; C, metaphase II). In comparison, *atm* exhibited a low level of chromosome fragmentation at normal temperature; interestingly, this fragmentation was significantly increased under heat stress (Fig. 5A, metaphase I; B, anaphase I; C, metaphase II). Examination of Arabidopsis with loss-of-function of the ATM ortholog protein ATR, however, did not reveal any defect in meiotic chromosome integrity at either temperature (Fig. 5A, metaphase I; B, anaphase I; C, metaphase II). Since heat stress reduced DSB formation in *atm*, the pronounced defects in chromosome integrity in *atm* at high temperature were likely attributable to impaired DSB repair.

**Figure 5.**
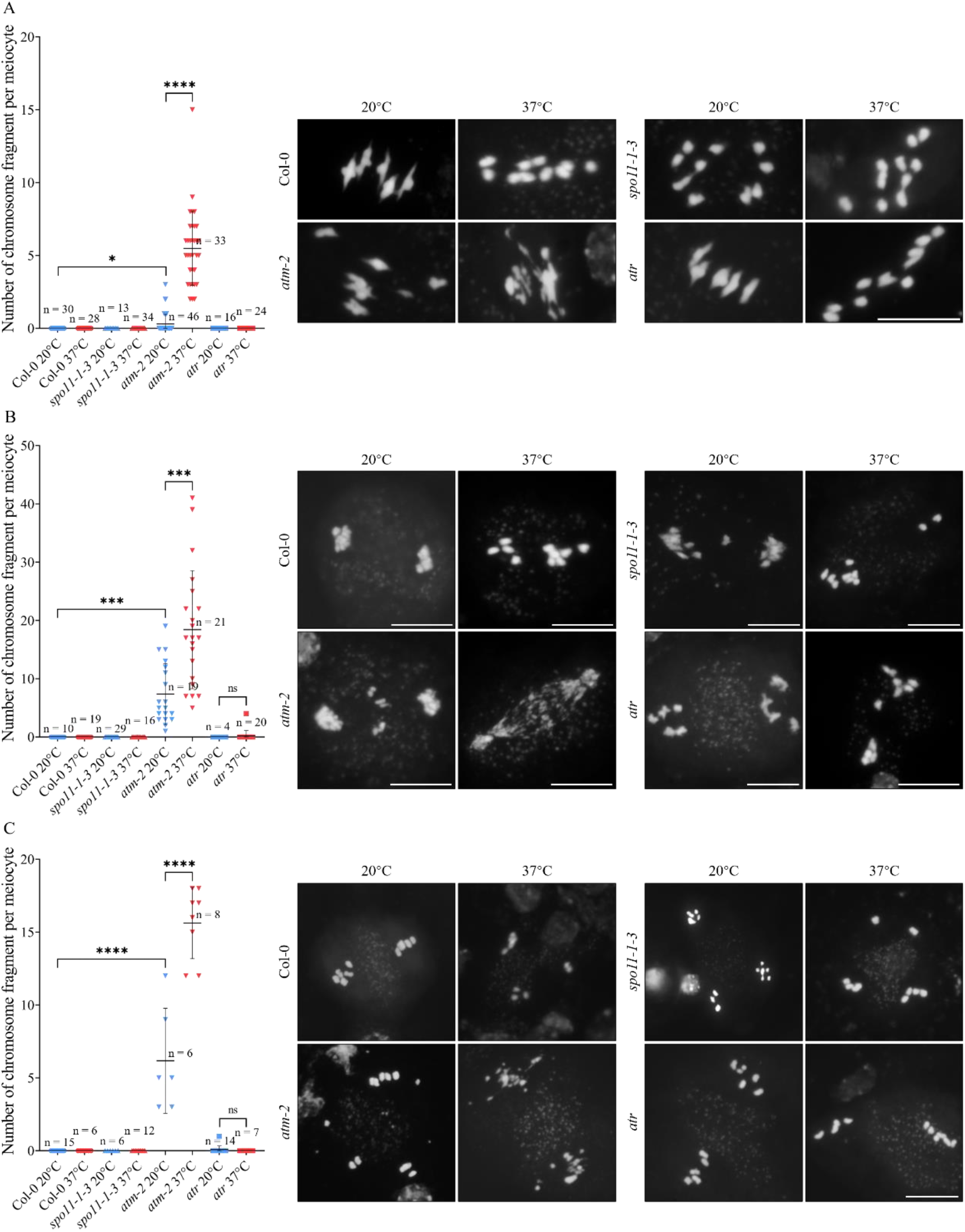
Analysis of chromosome spreads in Col-0, *spo11-1-3, atm-2* and *atr* at 20 and 37°C. A-C, Quantification of chromosome fragments in metaphase I (A), anaphase I (B) and metaphase II (C) meiocytes in Col-0, *spo11-1-3, atm-2* and *atr* at 20 and 37°C. Nonparametric and unpaired *t* tests were performed. n indicates the number of cells analyzed. **** indicates *P* < 0.0001; *** indicates *P* < 0.001;* indicates *P* < 0.05; ns indicates *P* > 0.05. Scale bars = 10 μm.

### Heat stress reduces chromosome fragmentation in *atm* in the *mre11* background

Upon DSB formation, the MRE11/RAD50/NBS1 (MRN) complex and ATM act in coordination to recognize and process DNA breaks and initiate DSB repair (Lavin, 2007; Paull, 2015). We thus wondered whether the essential function of ATM-mediated DSB repair at high temperature requires the MRN complex. To verify this, we took advantage of the *mre11* mutant, which is defective for dsDNA processing and DSB repair (Puizina et al., 2004; Šamanić et al., 2016; Samanić et al., 2013). At the control temperature, *mre11* showed significantly reduced ɤH2A.X foci formation (Supplemental Fig. S9), which was due to the impaired activation of ATM and the impaired H2A.X phosphorylation (Burma et al., 2001; Wang et al., 2019). We therefore did not use ɤH2A.X foci as the DSB indicator in *mre11* and instead evaluated the abundance of RAD51 and DMC1, as we did in *atm*. At 20°C, *mre11* and Col-0 showed similar numbers of RAD51 and DMC1 foci on zygotene chromosomes (Fig. 6A and B). This was unexpected, since it is thought that dysfunction of the MRN complex and resultant failure in single-stranded DNA generation compromise the loading efficiency of RAD51 and DMC1 on chromosomes (Bakr et al., 2015; Lukaszewicz et al., 2015; Mehnert et al., 2021). After heat stress treatment, both Col-0 and *mre11* showed a significantly reduced number of RAD51 and DMC1 foci (Fig. 6A and B), which manifested a decreased DSB abundance.

**Figure 6.**
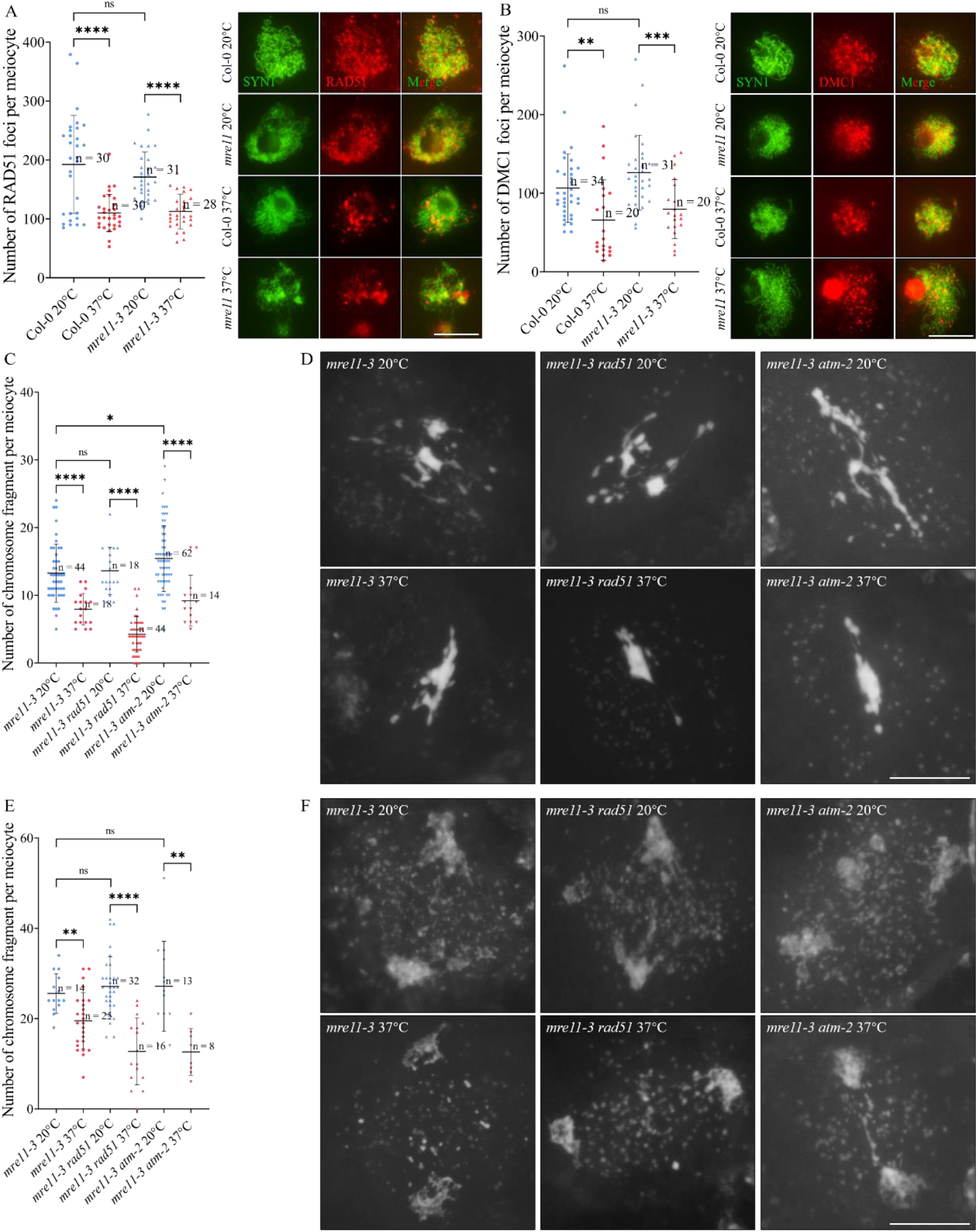
Heat stress reduces chromosome fragmentation in *mre11-3 atm-2*. A and B, Immunolocalization and quantification of RAD51 (A) and DMC1 (B) in Col-0 and *mre11-3* at 20 and 37°C. C and E, Quantification of chromosome fragments in metaphase I (C) and anaphase I meiocytes (E) in *mre11-3, mre11-3 rad51* and *mre11-3 atm-2* at 20 and 37°C. D and F, 4′,6-diamidino-2-phenylindole (DAPI)-staining of metaphase I (D) and anaphase I meiocytes (F) in *mre11-3, mre11-3 rad51* and *mre11-3 atm-2* at 20 and 37°C. Nonparametric and unpaired *t* tests were performed. n indicates the number of cells analyzed. **** indicates *P* < 0.0001; *** indicates *P* < 0.001; ** indicates *P* < 0.01; * indicates *P* < 0.05; ns indicates *P* > 0.05. Scale bars = 10 μm.

We next performed meiotic spread analysis and counted the number of chromosome fragments in *mre11*. At 20°C, metaphase I and anaphase I meiocytes in *mre11* showed an average of 13.27 and 25.57 fragments per meiocyte (Fig. 6C and D), which were decreased to 7.94 and 19.52, respectively, at 37°C (Fig. 6C and D). We introduced *spo11-1* into *mre11* and found that the impaired meiotic chromosome integrity, but not the mitosis defects, was suppressed in the double mutant (Supplemental Fig. S10). Overall, these data suggested that heat stress reduces DSB formation in *mre11*. On the other hand, at normal temperature, dysfunction of RAD51 in the *mre11* background did not alter the chromosome fragmentation level (Fig. 6C and D), which was in line with the fact that RAD51 acts downstream of the MRN complex in DSB repair. At 37°C, we observed a lowered abundance of fragments in *mre11 rad51* (Fig. 6C and D). To determine whether ATM-dependent chromosome integrity at high temperature is regulated in an MRN complex-mediated pathway, we introduced *atm* into the *mre11* background and analyzed the chromosome behaviors in the double mutant. At 20°C, we observed an elevated number of chromosome fragments in metaphase I meiocytes of *mre11 atm* compared with that in the single *mre11* mutant (Fig. 6C and D), which was probably due to the increase in DSB number caused by ATM dysfunction. After heat stress treatment, the number of fragments was significantly reduced in both metaphase I and anaphase I meiocytes of *mre11 atm* (Fig. 6C and D), indicating that the ATM-regulated DSB repair mechanism required for meiotic chromosome integrity is dependent on the normal function of the MRN complex.

Moreover, to determine whether ATM-dependent genome integrity at high temperature also exists in somatic cells, we analyzed mitotic chromosomes in somatic cells in unopened flower buds of Col-0, *atm* and *mre11*. Compared with Col-0, *mre11* exhibited a high level of abnormal chromosome interactions at anaphase (Supplemental Fig. S11), supporting the important role of MRE11 in the somatic DNA damage response (Puizina et al., 2004). Meanwhile, we consistently detected a low rate of irregular chromosome bridges in *atm* (Supplemental Fig. S11), suggesting that ATM plays a minor role in mitosis in Arabidopsis. After heat stress treatment, we did not observe any significant change in the level of mitosis defects in Col-0, *atm* or *mre11* (Supplemental Fig. S11), which suggested that the heat stress condition tested in our assay may not affect mitotic chromosome integrity in Arabidopsis flower tissues or that MRN-ATM signaling is not required for mitotic chromosome integrity at high temperature.

### Heat stress promotes chromosome integrity in *rad51* and *mnd1* relying on the presence of ATM

ATM activates and stabilizes RAD51 to facilitate DSB repair (Ahlskog et al., 2016; Yao et al., 2020). We thus questioned whether ATM-dependent chromosome integrity at high temperature is regulated through the RAD51-mediated pathway. To this end, chromosome integrity in the single *rad51* and double *rad51 atm* mutants was evaluated at control and high temperatures by counting the number of chromosome fragments. At 20°C, *rad51* harbored an average of 3.86 and 20.08 chromosome fragments per metaphase I and anaphase I meiocytes, respectively; comparatively, *rad51 atm* showed a higher level of chromosome fragmentation owing to the *atm*-induced increase in DSB number (8.43 at metaphase I; 27.16 at anaphase I) (Fig. 7A and B). At high temperature, in line with what we previously reported, the number of chromosome fragments in *rad51* was lowered to 1.05 and 7.22 per meiocyte at metaphase I and anaphase I, respectively (Fig. 7A and B) (Ning et al., 2021). Interestingly, *rad51 atm* retained a high level of chromosome instability after heat stress treatment, showing an average of 7.45 and 30.38 fragments per metaphase I and anaphase I meiocyte, respectively; these values were not significantly different from the corresponding numbers of fragments at 20°C (Fig. 7A and B). To confirm that heat stress reduced DSB formation in *rad51 atm*, we compared the abundance of DMC1 foci in zygotene meiocytes of Col-0 and *rad51 atm* (Fig. 7C). At 20°C, *rad51 atm* exhibited fewer DMC1 foci than Col-0, likely due to the impairment of RAD51 function and the resultant interfered DMC1 chromosome localization (Chen et al., 2021). At 37°C, the number of DMC1 foci in Col-0 and *rad51 atm* was lowered, indicating reduced DSB formation. Therefore, these findings demonstrated that heat stress does not rescue chromosome integrity in *rad51* when ATM is absent.

**Figure 7.**
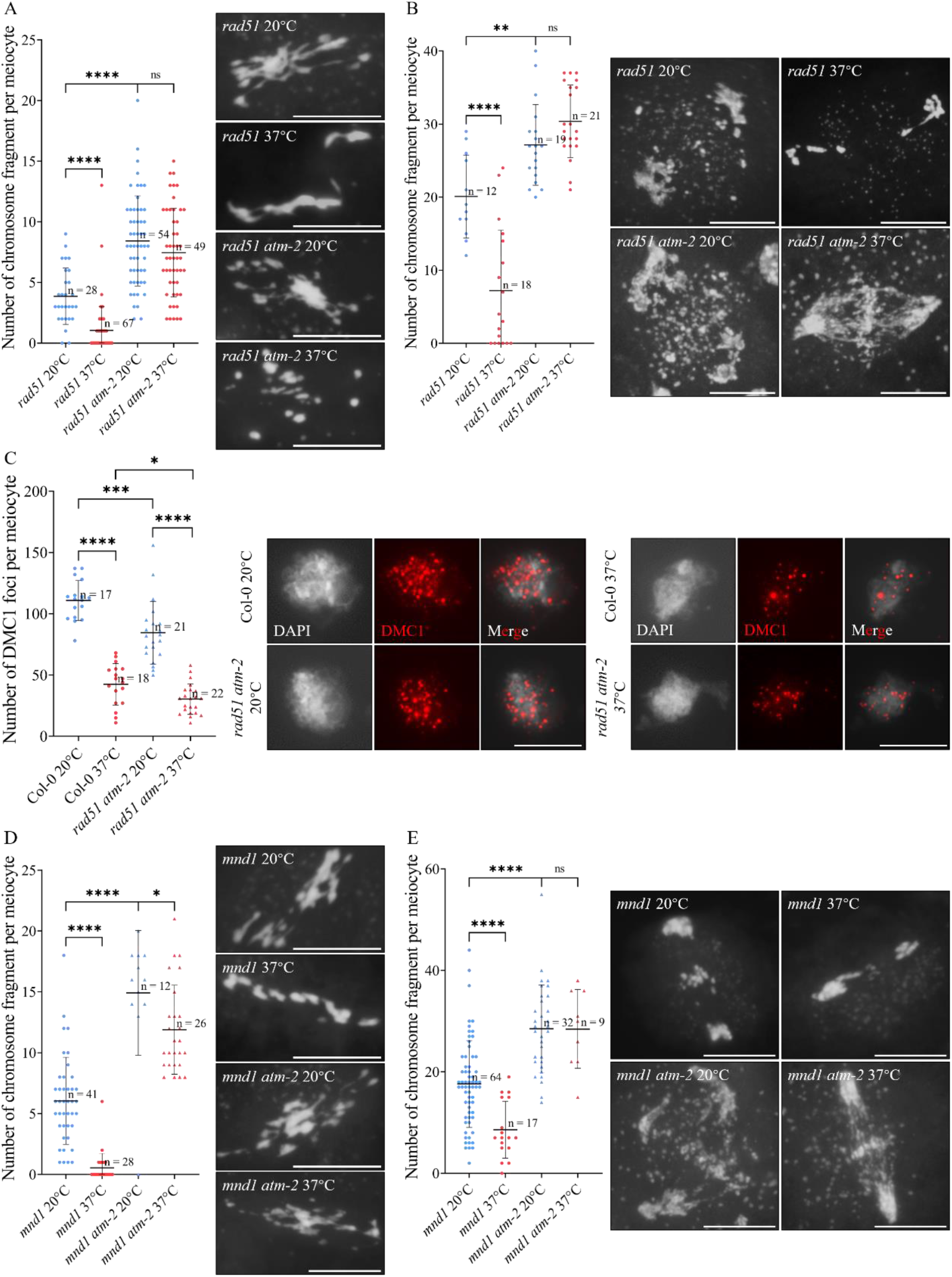
Heat stress reduces DSB formation but does not rescue chromosome fragmentation in the *rad51* and *mnd1* mutants in the absence of ATM. A and B, Quantification of chromosome fragments in metaphase I (A) and anaphase I meiocytes (B) in the *rad51* and *rad51 atm-2* mutants at 20 and 37°C. C, Immunolocalization and quantification of DMC1 on zygotene chromosomes in the *rad51 atm-2* mutant at 20 and 37°C. D and E, Quantification of chromosome fragments in metaphase I (D) and anaphase I meiocytes (E) in the *mnd1* and *mnd1 atm-2* mutants at 20 and 37°C. Nonparametric and unpaired *t* tests were performed. n indicates the number of cells analyzed. **** indicates *P* < 0.0001; *** indicates *P* < 0.001; ** indicates *P* < 0.01; * indicates *P* < 0.05. Scale bars = 10 μm.

To confirm that ATM-mediated DSB repair at high temperature is regulated through a pathway parallel to the RAD51-mediated one, we analyzed chromosome integrity in Arabidopsis plants with dysfunction of MEIOTIC NUCLEAR DIVISION 1 (MND1), which acts together with HOMOLOGOUS-PAIRING PROTEIN 2 (HOP2) to facilitate single-stranded DNA loading and the strand exchange activity of RAD51 and DMC1 during DSB repair and HR (Pezza et al., 2007; Uanschou et al., 2013; Vignard et al., 2007; Zhao et al., 2014). At 20°C, *mnd1* showed an average of 6.05 and 17.61 chromosome fragments per meiocyte at metaphase I and anaphase I, respectively, which were elevated to 14.92 and 28.50 when combined with the ATM loss-of-function mutation (Fig. 7D and E). After heat stress treatment, chromosome integrity in the metaphase I and anaphase I meiocytes of *mnd1* was largely rescued (Fig. 7D and E). In contrast, in *mnd1 atm*, although we detected a significant decrease in chromosome fragment number in metaphase I meiocytes after heat stress treatment (Fig. 7D; 14.92 at 20°C; 11.88 at 37°C), there was no difference in the number of chromosome fragments in anaphase I meiocytes between plants at the two temperatures (Fig. 7E; 28.50 at 20°C; 28.44 at 37°C). Collectively, these data suggested that ATM-dependent chromosome stability at extreme high temperature is regulated at least partially through a RAD51-independent DSB repair pathway.

### Heat stress further reduces DMC1 localization in the RAD51-SMC5/6 signaling mutants

RAD51 facilitates DMC1-mediated DSB repair by suppressing the negative impact of the SMC5/6 complex (Chen et al., 2021). The further reduction of DMC1 foci in *rad51 atm* at high temperature suggested that high temperature reduces the chromosome localization of DMC1 independently of RAD51 (Fig. 7C). To verify this, we compared the number of DMC1 foci on zygotene chromosomes in a series of RAD51-SMC5/6 signaling mutants at 20 and 37°C. We took advantage of the single *sni1* and *asap1*, and the double *rad51 sni1* and *rad51 asap1* mutants which have been characterized previously (Chen et al., 2021). At 20°C, *rad51* exhibited a significantly lower number of DMC1 foci than Col-0, whereas *sni1* and *asap1* showed increased DMC1 accumulation (Fig. 8A, B, D, F and H). The double *rad51 sni1* and *rad51 asap1* mutants demonstrated partially compromised DMC1 localization compared with the single *sni1* and *asap1* mutants (Fig. 8A, J and L). These observations supported the proposed function of RAD51-SMC5/6 signaling in regulating DMC1 (Chen et al., 2021). At 37°C, as in Col-0, the number of DMC1 foci was reduced in all the mutants (Fig. 8A, C, E, G, I, K and M). Taken together, these findings support the model that heat stress reduces DMC1 accumulation on chromosomes independently of RAD51-SMC5/6 signaling, possibly via a direct impact on DSB formation.

**Figure 8.**
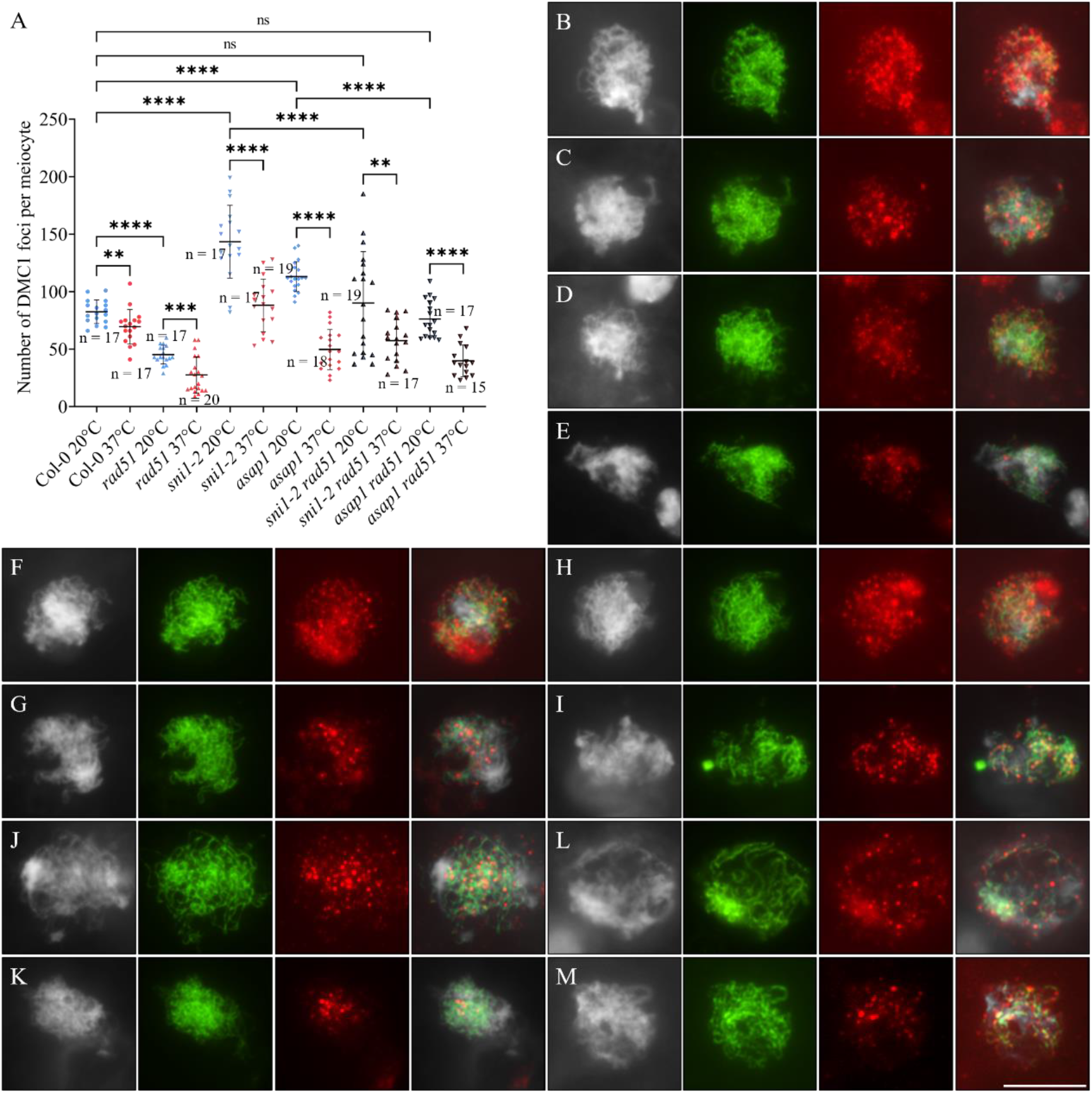
Heat stress reduces the chromosome localization of DMC1 independent of RAD51-SMC5/6 signaling. A, Graph showing the quantification of DMC1 foci on zygotene chromosomes of Col-0, *rad51, sni1-2, asap1, sni1-2 rad51* and *asap1 rad51* plants. Nonparametric and unpaired *t* tests were performed. n indicates the number of cells analyzed. **** indicates *P* < 0.0001; *** indicates *P* < 0.001; ** indicates *P* < 0.01; ns indicates *P* > 0.05. B-M, Dual-immunolocalization of SYN1 and DMC1 on zygotene chromosomes of Col-0 (B and C), *rad51* (D and E), *sni1-2* (F and G), *asap1* (H and I), *sni1-2 rad51* (J and K) and *asap1 rad51* (L and M) plants at 20 (B, D, F, H, J and L) and 37°C (C, E, G, I, K and M). White, DAPI; green, SYN1; red, DMC1. Scale bar = 10 μm.

### Heat stress reduces chromosome fragmentation in *atm* in the *syn1* background

In Arabidopsis, the chromosome axis plays multiple roles during HR, including roles in DSB formation, homolog synapsis and CO formation (Lambing et al., 2022; Lambing et al., 2020). To explore the potential role of the chromosome axis in ATM-mediated DSB repair at high temperature, we performed chromosome spread analysis and counted the chromosome fragments in metaphase I and anaphase I meiocytes in the single *syn1* and double *syn1 atm* mutants at 20 and 37°C (Fig. 9). The *syn1* mutant has a defect in DSB repair and consequent defect in meiotic chromosome integrity; in our assay, this mutant showed an average of 3.53 and 19.60 chromosome fragments per meiocyte at metaphase I and anaphase I, respectively, at 20°C (Fig. 9A and B). The *atm* mutation did not alter the fragmentation level in *syn1* (Fig. 9A and B), probably due to a defect in DSB formation caused by SYN1 dysfunction (Lambing et al., 2020). After heat stress treatment, because of the reduced DSB formation, the number of chromosome fragments in *syn1* was lowered to 1.66 and 14.07 per meiocyte at metaphase I and anaphase I, respectively (Fig. 9A and B) (Ning et al., 2021). Interestingly, heat-induced reduction of chromosome fragments also occurred in *syn1 atm* (Fig. 9A and B), suggesting that high temperature did not further damage chromosome integrity in *atm* when SYN1 was absent. Overall, these data suggested that ATM-mediated DSB repair at high temperature is dependent on SYN1.

**Figure 9.**
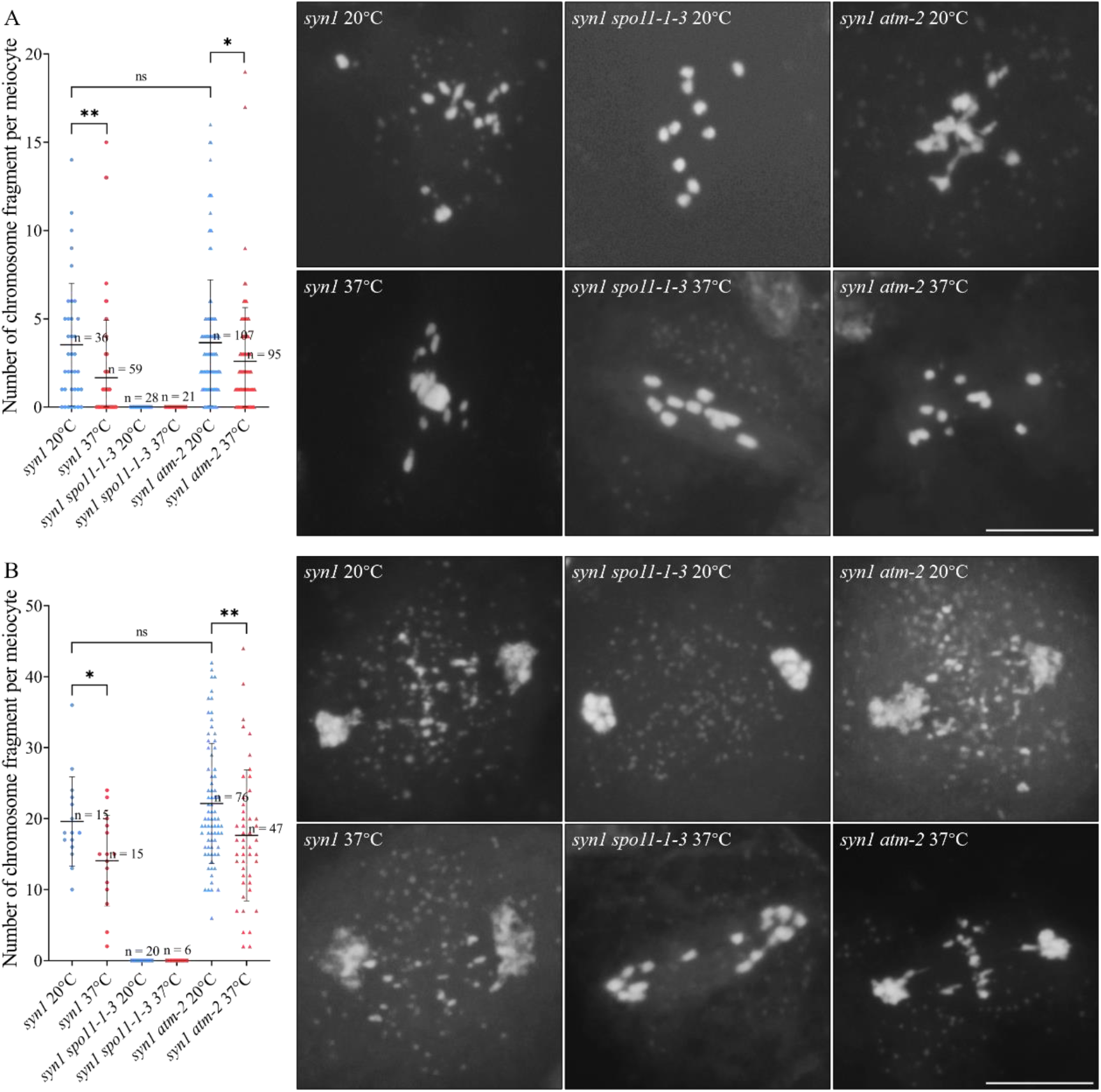
Chromosome spread analysis in chromosome axis-related mutants. A and B, Quantification of chromosome fragments in metaphase I (A) and anaphase I (B) meiocytes in the *syn1, syn1 spo11-1-3* and *syn1 atm-2* mutants at 20 and 37°C. Nonparametric and unpaired *t* tests were performed. n indicates the number of cells analyzed. ** indicates *P* < 0.01; * indicates *P* < 0.05. ns indicates *P* > 0.05. Scale bars = 10 μm.

### ATM dysfunction does not alter the impact of heat stress on SC and CO formation

At extremely high temperature, the formation of the synaptonemal complex (SC) and COs in wild-type Arabidopsis is impaired (Ning et al., 2021). To test whether ATM plays a role in these processes, we analyzed homolog synapsis and CO formation in *atm* at 20 and 37°C. The morphology of the lateral element of SC was examined by performing dual-immunolocalization of the ASY1 and ASY4 proteins, both of which are required for normal synapsis and are sensitive to heat stress (Armstrong et al., 2002; Chambon et al., 2018; Fu et al., 2021; Ning et al., 2021). At normal temperature, ASY1 and ASY4 signals in *atm* and *mre11* revealed a linear configuration and were fully overlapped on the whole region of zygotene chromosomes, similar to the case in Col-0; meanwhile, at 37°C, dotted ASY1 and ASY4 signals were observed in Col-0 and the *atm* and *mre11* mutants (Fig. 10A). These findings suggested that the impaired DSB repair at early prophase I stages does not interfere with the formation and the heat response of the lateral element of SC. Meiotic spread analysis revealed normal synapsis of homologous chromosomes at pachytene in *atm* at 20°C, which, however, was completely abolished at 37°C (Fig. 10B and C). Dual-immunolocalization of SYN1 and ZYP1 confirmed the disruption of SC formation in heat stressed *atm*, in which the linear configuration of ZYP1 was disrupted, becoming fragmentary and/or dotted (Fig. 10B and C). In comparison, *mre11* displayed damaged homolog synapsis and SC formation at both temperatures (Supplemental Fig. S12).

**Figure 10.**
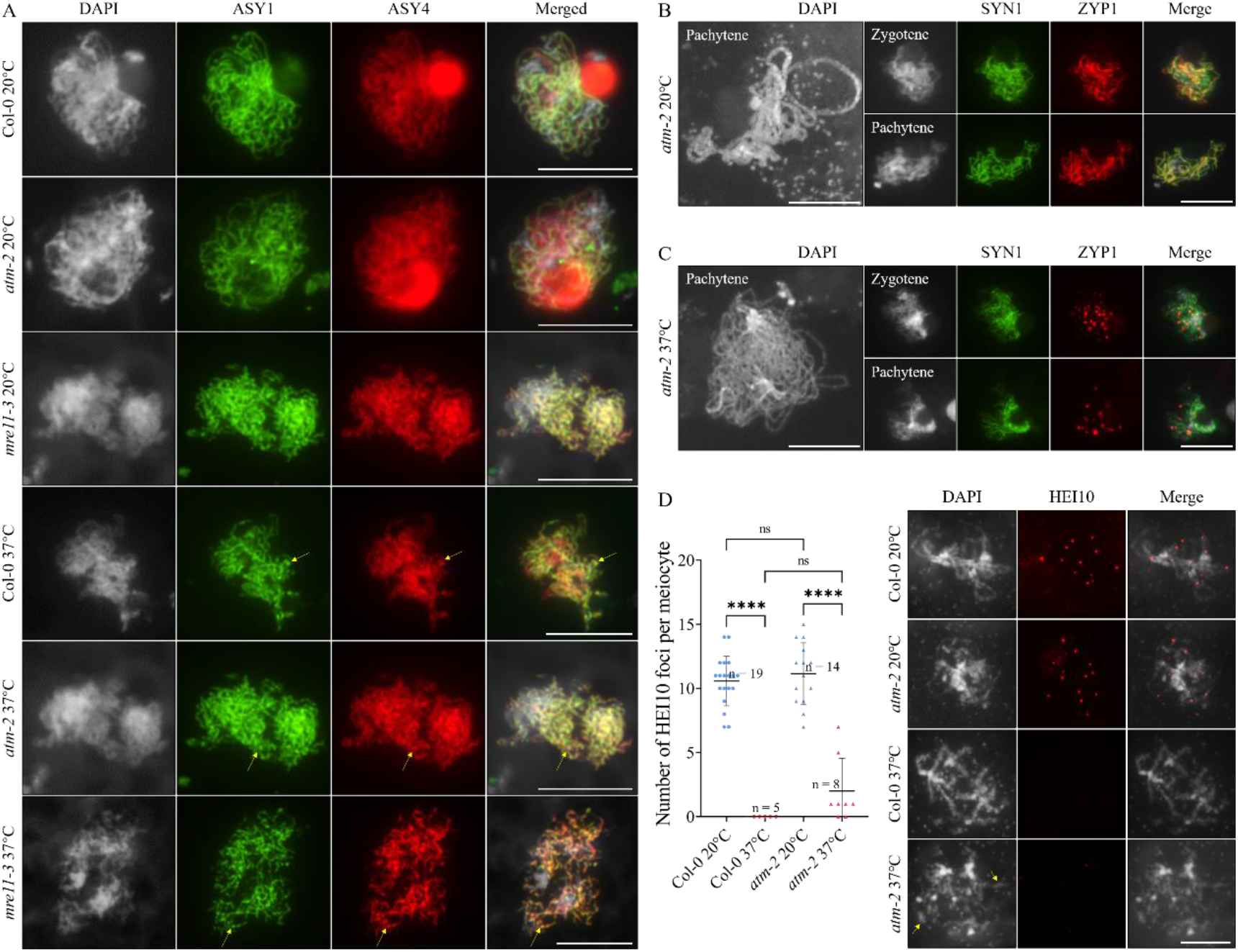
Analysis of SC and CO formation in *atm-2* at 20 and 37°C. A, Dual-immunolocalization of ASY1 and ASY4 in Col-0, *atm-2* and *mre11-3* at 20 and 37°C. Yellow arrows indicate the dotted configuration of ASY1 and/or ASY4. B and C, DAPI-staining of pachytene chromosomes, and dual-immunolocalization of SYN1 and ZYP1 on zygotene- and pachytene chromosomes in *atm-2* at 20 (B) and 37°C (C). D, Immunolocalization and quantification of HEI10 on diplotene chromosomes in Col-0 and *atm-2* at 20 and 37°C. Yellow arrows indicate the fragmented chromosomes. Nonparametric and unpaired *t* tests were performed. n indicates the number of cells analyzed. **** indicates *P* < 0.0001; ns indicates *P* > 0.05. Scale bars = 10 μm.

CO formation in *atm* was evaluated by performing immunolocalization of HEI10 protein, a marker of class I-type COs (Chelysheva et al., 2012; Higgins et al., 2004; Wang et al., 2012; Ziolkowski et al., 2017). At 20°C, diplotene meiocytes in *atm* showed a similar number of HEI10 foci as Col-0 (Fig. 10D; 10.58, Col-0; 11.14, *atm*), which supported that dysfunction of ATM does not affect class I-type CO formation in Arabidopsis grown at normal temperatures (Kurzbauer et al., 2021). At 37°C, intact but unsynapsed univalent chromosomes in Col-0 exhibited impaired HEI10 localization (Fig. 10D). In *atm*, although we occasionally detected a small number of HEI10 foci on fragmented chromosome bodies in some meiocytes, there was no significant difference in HEI10 pattern compared with that in heat-stressed Col-0 (Fig. 10D). Collectively, these data demonstrated that ATM dysfunction does not influence the impact of heat stress on SC and CO formation in Arabidopsis.

## Discussion

### Heat stress reduces DSB formation possibly by interfering with the chromosome localization of DSB formation proteins

We previously reported that extreme high temperature (37°C) reduces the generation of DNA double-strand breaks (DSBs) in *Arabidopsis thaliana* (Ning et al., 2021). In this study, we showed that the chromosome localization of SPO11-1 and DFO, which are key DSB formation proteins (Grelon et al., 2001; Hartung et al., 2007; Stacey et al., 2006; Zhang et al., 2012), was compromised at high temperature (Fig. 1B and Fig. 2B). In the proposed model of the DSB formation complex, the amounts of different DSB formation proteins need to be coordinately regulated; meanwhile, subcomplexes of specific proteins are formed by direct and/or indirect interactions (Tang et al., 2017; Vrielynck et al., 2021). The reduced number of SPO11-1 and DFO proteins on the chromosomes may interfere with organization and/or the structure of the subcomplex or the protein interaction environment within the DSB formation machinery, resulting in perturbed function (Vrielynck et al., 2021). We found that the localization of DFO was interfered in *spo11-1* meiocytes and that heat stress reduced number of DFO foci in a manner dependent on the presence of SPO11-1 (Fig. 2C). It is possible that the interfered localization of DFO was at least partially a secondary effect of high temperature on the chromosome loading of SPO11-1. In the DSB complex, MTOPVIB and its interacting partner PRD1 act as the core factors to establish a platform that facilitates DSB formation protein associations through specific protein-protein interactions. MTOPVIB channels the connections between SPO11-1 (and/or -2) and the DFO/PRD2/PHS1 subcomplex (Tang et al., 2017; Vrielynck et al., 2021). We speculate that the chromosome localization of MTOPVIB and PRD1 may also be affected by heat stress. Determination of the chromosome localization patterns of other DSB formation proteins at increased temperature could contribute to the understanding how heat stress influence structure and/or function of the DSB complex.

### ATM does not mediate the reduced DSB formation at high temperature

In Arabidopsis plants with ATM dysfunction, in accordance with the increased number of DSBs formed (Fig. 3) (Kurzbauer et al., 2021; Yao et al., 2020), we found that the number of SPO11-1 and DFO foci was also increased (Fig. 4). These findings hint that in Arabidopsis, one of the mechanisms by which ATM restricts DSB formation may be by limiting the accumulation and/or chromosome localization of DSB formation proteins (Carballo et al., 2013; Garcia et al., 2015; Kurzbauer et al., 2021; Lange et al., 2011; Paiano et al., 2020). We considered the possibility that ATM mediates the reduced DSB formation at high temperature, because: (1) ATM negatively regulates meiotic DSB formation (Carballo et al., 2013; Kurzbauer et al., 2021; Lange et al., 2011); (2) the expression of ATM is upregulated at high temperature (Supplement Fig. S6B) (Ning et al., 2021); and (3) both heat stress and ATM dysfunction reduce the chromosome localization of SPO11-1 and DFO (Fig. 1B; Fig. 2B). However, this hypothesis was undermined by the observations that ATM dysfunction did not affect the compromised chromosome localization of RAD51, DMC1, SPO11-1 and DFO proteins at high temperature (Fig. 3; Fig. 4). Therefore, our data can conclude that ATM does not mediate the heat-induced reduction in DSB formation in Arabidopsis. On the other hand, at normal temperature, Arabidopsis plants with impaired ATM function have a defect in meiotic chromosome integrity (Fig. 5) (Kurzbauer et al., 2021; Yao et al., 2020), which could be caused by increased DSB abundance or impacted DSB repair that likely occurs post RAD51- and DMC1-nucleofilaments formation, or both. It has been reported that elevated temperature can promote progressing of early prophase in Arabidopsis meiosis, but this effect is compromised when ATM is absent (De Jaeger-Braet et al., 2022). Therefore, increased accumulation of RAD51 and DMC1 protein foci on *atm* early prophase chromosomes may result from delayed and/or perturbed DSB repair (Fig. 3) (Kurzbauer et al., 2021; Yao et al., 2020). Here, we showed that the number of chromosome fragments in *atm* meiocytes was increased at extreme high temperature (Fig. 5). Since heat stress reduced DSB formation in *atm*, the pronounced chromosome fragments in *atm* at high temperature (Fig. 5) plausibly result from a further attenuation of DSB repair ability.

### ATM-mediated DSB repair is required for meiotic genome stability at high temperature

ATM, whose activation is stress inducible, plays a central role, together with the MRN complex, in the sensing and repair of DSBs using diverse strategies (Shibata and Jeggo, 2021; Williams and Zhang, 2021). We showed that, compared with that in *atm*, the chromosome fragmentation phenotypes in both *mre11, mre11 rad51* and *mre11 atm* could be partially complemented at high temperature (Fig. 6C-F), only manifesting a reduction in DSB formation upon heat stress. Hence, we speculate that ATM-mediated DSB repair at high temperature relies on the function of MRE11 and occurs after double-stranded DNA end resection, which supports a master role of the MRN complex in regulating DSB repair (Kanaar and Wyman, 2008; Puizina et al., 2004; Williams et al., 2010; Williams et al., 2007).

In mammals, ATM and MRE11 process DSB ends to generate single-stranded DNAs (ssDNAs), which facilitates the recruitment of the ATM ortholog ATR for subsequent DSB repair events. This mechanism specifically takes place in a cell cycle-dependent manner (Jazayeri et al., 2006). We showed here that ATR dysfunction did not induce any obvious defect in meiotic chromosome integrity at normal or high temperatures (Fig. 5), suggesting that the ATM-to ATR-mediated DSB repair machinery switching mechanism is not involved in this context (Shiotani and Zou, 2009). Additionally, it is possible that ATR may play a predominant role in mitotic but not meiotic DSB repair in plants (Amiard et al., 2010; Amiard et al., 2011).

ATM regulates DSB repair primarily through the homologous recombination (HR)-dependent pathway (Williams and Zhang, 2021). In mammals, ATM facilitates HR-dependent DSB repair by promoting DSB end processing and antagonizing the assembly of proteins involved in error-prone repair pathways, including nonhomologous end joining (NHEJ), which, in bacteria, is activated upon heat shock (Britton et al., 2020; Dupuy et al., 2018; Muraki et al., 2013). Meanwhile, it has recently been reported in *Caenorhabditis elegans* that ATM/ATR maintains the bias of meiotic DSB repair pathway choice (Láscarez-Lagunas et al., 2022). Specifically, ATM/ATR phosphorylates SYP-4 protein, a central element of SC, at the S447 site upon DSB formation, which prevents meiotic DSBs to be repaired through the NHEJ pathway (Láscarez-Lagunas et al., 2022). In plants, not much is known about the role of the HNEJ pathway in meiotic DSB repair especially under stressful conditions. The possibility cannot be excluded that in Arabidopsis meiocytes, dysfunction of ATM may perturb the HR-dependent DSB repair pathway and evoke an ectopic activation of NHEJ signaling that might be less-efficient at extremely high temperatures, which consequently leads to impaired genome integrity.

A large-scale proteomic analysis in mammals revealed that ATM controls a very complex DSB damage response network (Matsuoka et al., 2007); however, the substrates of ATM in plants, especially in sexual mother cells, are largely unclear. We showed here that the heat-induced decrease in chromosome fragmentation in the *rad51* and *mnd1* mutants was abolished when ATM was absent (Fig. 7), suggesting that a further disruption of DSB repair caused by ATM dysfunction compromised the effect of heat stress on DSB reduction in *rad51* and *mnd1*. We propose that ATM may control a DSB repair pathway that occurs parallel to RAD51 and is required for meiotic chromosome integrity at high temperature. As such, XRCC3, an ATM-targeted RAD51 paralog protein that plays a conserved role in facilitating DSB repair by promoting RAD51 chromosome loading and the processing of homologous recombination intermediates (Bishop et al., 1998; Brenneman et al., 2002; Pierce et al., 1999; Su et al., 2017; Takata et al., 2001; Tang et al., 2014; Zhang et al., 2015), may not be a candidate.

On the other hand, meiotic recombination characteristics, including DSB and CO formation and distribution, are highly correlated with chromatin features that are both diverse and divergent among species (Underwood et al., 2018; Zelkowski et al., 2019). For example, in Arabidopsis, DSBs are less enriched at heterochromatin regions (Lambing et al., 2020). Analysis of genome-wide high-resolution chromatin configuration in Arabidopsis seedlings has revealed that heat stress induces global reorganization of chromatin structures and promotes heterochromatin decondensation (Sun et al., 2020). However, it is not yet clear how heat stress influences chromatin features and dynamics in plant meiocytes. In mammals, ATM has been shown to specifically facilitate the repair of DSBs within heterochromatin regions, where it promotes the access ability of DSB repair proteins by phosphorylating heterochromatin factors and the associated remodeling of chromatin (Goodarzi et al., 2010; Goodarzi et al., 2011; Goodarzi et al., 2008). Considering the conserved function of ATM, such an ATM-regulated chromatin status-dependent DSB repair mechanism might also exist in plants. It might be possible that extreme high temperature alters the proportion and/or the repair efficiency of the DSBs programmed to be processed by ATM by influencing chromosome organization at early meiotic recombination stages.

### ATM-mediated DSB repair relies on functional chromosome axis

In the *syn1* mutant, we showed that ATM dysfunction did not induce an increased chromosome fragmentation level (Fig. 9). This could be caused by the interfered DSB formation due to the disorganization of the chromosome axis (Lambing et al., 2022; Lambing et al., 2020). Compared with its function in ensuring normal DSB formation, SYN1 is more required for DSB repair, since under normal growth conditions, *syn1* shows a severe chromosome fragmentation phenotype that can be fully rescued in the *spo11-1* background (Fig. 9) (Bai et al., 1999). Interestingly, we found that dysfunction of SYN1 contributed to increased meiotic chromosome integrity in *atm* at high temperature (Fig. 9), manifesting a prominent effect of reduced DSB formation. We speculate that heat stress may not be able to further perturb the DSB repair ability in *syn1 atm*. It is possible that ATM-dependent DSB repair relies on the normal organization and/or function of the SYN1-mediated axis, which establishes a platform for the localization and/or function of other recombination proteins (Lambing et al., 2022; Shahid, 2020; Vrielynck et al., 2021). The roles of the chromosome axis in meiotic DSB repair under both normal and stress conditions await further investigation. Taken together, our study suggests that ATM regulates a DSB repair mechanism that acts at least partially in a RAD51-independent manner but relies on functional SYN1-mediated chromosome axis, which protects meiotic genome stability in plants at extreme high temperature.

## Material and Methods

### Plant materials and growth conditions

*Arabidopsis thaliana* wild-type Columbia-0 (Col-0), *spo11-1-3* (SALK_146172) (Xue et al., 2018), *rad51* (GABI_134A01) (Li et al., 2004), *atm-2* (SALK_006953) (Kurzbauer et al., 2021), *atr* (SALK_032841) (Wang et al., 2021), *mre11-3* (SALK_054418) (Puizina et al., 2004), *mnd1* (SALK_110052) (Vignard et al., 2007), *asap1* (GABI_218F01) (Chen et al., 2021), *sni1-2* (SAIL_298_H07) (Chen et al., 2021), *asap1 rad51* (Chen et al., 2021), *sni1-2 rad51* (Chen et al., 2021), *dfo-1* (Zhang et al., 2012) and *syn1* (SALK_137095) (Wang et al., 2020) mutants were used in this study. The double mutants were generated by crossing the corresponding single heterozygous mutants. The *atm-2* plants expressing *pSPO11-1::SPO11-1-GFP* were generated by crossing the Col-0 plants expressing *pSPO11-1::SPO11-1-GFP* with the *atm-2* mutant using the former one as a pollen donor. Genotyping was performed by polymerase chain reaction (PCR) using the primers listed in Supplemental Table S1. Seeds were germinated in K1 medium for 6-8 days and seedlings were transferred to soil and cultivated in a growth chamber with a 16 h day/8 h night, 20°C, and 50% humidity.

### High temperature treatment

Young flowering Arabidopsis plants were treated with 37°C for 24 h in a humid chamber under a 16 h day/8 h night period. All the treatment assays were initiated from 8:00-10:00 AM. Flower samples were collected or fixed upon the completion of the treatment.

### Generation of the *pSPO11-1::SPO11-1-GFP* reporter

To generate the *pSPO11-1::SPO11-1-GFP* reporter, a 3970 bp genomic sequence of SPO11-1 was amplified by PCR using primers (SPO11-1-attB1-F and SPO11-1-attB2-R) flanked with attB sites. The PCR fragment was then integrated into the *pDONR221* vector via Gateway BP reaction. Next, a SmaI restriction site was introduced in front of the stop codon of SPO11-1 by PCR. The resulting construct was then linearized by SmaI restriction and ligated with the GFP fragment, followed by Gateway LR reaction with the destination vector *pGWB501*.

### Meiotic chromosome spread analysis

Inflorescences of young, flowering Arabidopsis plants were collected and fixed in pre-cooled Carnoy’s fixative for at least 24 h. Flower buds beyond the tetrad stage were removed, and the remaining buds were washed twice with distilled water and once with citrate buffer (10 mM, pH = 4.5), followed by incubation in a digestion enzyme mixture (0.3% pectolyase and 0.3% cellulase in citrate buffer, 10 mM, pH = 4.5) at 37°C for 2.5 h. Digested flower buds were subsequently washed once in distilled water and macerated in 3 μL distilled water on a glass slide. Two aliquots of 10 μL 60% acetic acid were added to the slide, which was then placed on a hotplate at 45°C for 1.5 min. When the slide was dried, it was flooded with 200 μL freshly-prepared ice-cold Carnoy’s fixative and subsequently air dried. Eight microliters of 4′,6-diamidino-2-phenylindole (DAPI) (5 μg/mL) in Vectashield antifade mounting medium was added to the slide, and the coverslip was mounted and sealed with nail polish. Slides were immediately examined under a fluorescence microscope, or stored at -20°C. The slides were examined for both meiotic and mitotic cells (from somatic tissue within the flower buds).

### Generation of antibodies

The anti-AtHEI10 antibody was generated in rabbits against an 80 amino acid peptide of AtHEI10 conjugated to KLH. The anti-AtDFO antibody was generated in mice against a 25 amino acid peptide of AtDFO conjugated to KLH. The anti-AtRAD51 antibody was generated in mice using the full-length AtRAD51 protein as an antigen, as previously reported (Mercier et al., 2003).

### Protein immunolocalization

Immunolocalization was performed by referring to (Chelysheva et al., 2010; Wang et al., 2014). Antibody against SYN1 (rabbit) (Ning et al., 2021) were diluted 1:500; antibody against AtSYN1 (mouse) (Fu et al., 2021) was diluted 1:200; antibodies against AtRAD51 (mouse), ASY1 (mouse) (Fu et al., 2021), ASY4 (rabbit) (Fu et al., 2021) and AtHEI10 (rabbit) were diluted 1:150; and antibodies against ɤH2A.X (rabbit) (Fu et al., 2021), AtZYP1 (rabbit) (Ning et al., 2021), AtDMC1 (rabbit) (Ning et al., 2021), AtDFO (mouse) and GFP (rabbit) (Invitrogen, PA1-980A) were diluted 1:100. The secondary antibodies goat anti-rabbit IgG (H+L) cross-adsorbed secondary antibody Alexa Fluor 555 (Invitrogen, A-32732), goat anti-rabbit IgG (H+L) highly cross-adsorbed secondary antibody Alexa Fluor Plus 488 (Invitrogen, A-32731), goat anti-mouse IgG (H+L) highly cross-adsorbed secondary antibody Alexa Fluor Plus 488 (Invitrogen, A32723) and F(ab’)2-goat anti-mouse IgG (H+L) cross-adsorbed secondary antibody Alexa Fluor Plus 555 (Invitrogen, A48287) were diluted to 10 μg/mL. The slides were immediately examined under a fluorescence microscope or stored at -20°C.

### Quantification of fluorescent foci and chromosome fragments

Counting of fluorescent foci was performed as previously reported (Fu et al., 2021; Ning et al., 2021). In brief, ImageJ software was used to merge the DAPI- and/or GFP- and RFP-channel images, and only the fluorescent foci that coincided with DAPI-stained and/or GFP-labeled chromosomes were quantified. Quantification of chromosome fragments was performed as described previously (Kurzbauer et al., 2021). In brief, DAPI-stained chromosome bodies that were obviously brighter than organelles were counted, followed by the subtraction of the number of expected chromosomes (5 at metaphase I, 10 at anaphase I and 20 at metaphase II).

### Quantitative reverse transcription PCR (RT-qPCR)

The quantitative reverse transcription PCR (RT-qPCR) used to study the gene expression level was performed as previously described with some minor modifications (Ning et al., 2021). Total RNA was isolated from meiosis-staged flower buds of Arabidopsis Col-0 plants grown at 20°C, or plants stressed at 37°C for 24 h. *TAP42 Interacting Protein of 41 kDA* (*TIP41*) was used as the reference gene (Škiljaica et al., 2022). The relative expression fold-change is shown in the figures. The primers used are listed in Supplemental Table S1.

### Microscope and image adjustment

Fluorescence images were captured by an Olympus IX83 inverted fluorescence microscope with an X-Cite lamp and a Prime BSI camera. Dual fluorescence images and Z-stacks were processed using ImageJ. The brightness and contrast settings of the images were adjusted using PowerPoint 2016.

### Statistical analysis

Significance analysis was performed using nonparametric and unpaired *t* tests with GraphPad Prism software (version 8). The number of cells analyzed or the number of biological replicates is shown in the figures and/or legends.

## Author contributions

J.Z. performed most of the laboratory experiments; X.G. analyzed chromosome spread and HEI10 localization; Z.R. analyzed gene expression and performed statistical analysis; H.F. analyzed DMC1 localization in the SMC5/6 complex mutants and contributed to meiotic spread analysis; C.Y. generated the *spo11-1-3*^*pSPO11-1::SPO11-1-GFP*^ transgenic line and analyzed the data; Q.L. contributed to gene expression analysis; M.Z. analyzed the *dfo* meiotic spread; W.W., C.W. and A.S. contributed to the data analysis and edited the manuscript; B.L. conceived the project, analyzed the data, and wrote and edited the manuscript. All the authors have read and agreed with the submission of the manuscript.

### Accession numbers

Accession numbers of genes studied in this work: *ATM* (AT3G48190), *ATR* (AT5G40820), *SPO11-1* (AT3G13170), *DFO* (AT1G07060), *MRE11* (AT5G54260), *RAD51* (AT5G20850), *DMC1* (AT3G22880), *ASY1* (AT1G67370), *ASY4* (AT2G33793), *ZYP1A* (AT1G22260), *SYN1* (AT5G05490), *HEI10* (AT1G53490), *SNI1* (AT4G18470), *ASAP1* (AT2G28130), *HSFA1a* (AT4G17750) and *TIP41* (AT4G34270).

## Supplemental data

Supplemental Figure S1. Phenotyping of *spo11-1-3* mutant plants expressing the *pSPO11-1::SPO11-1-GFP* reporter.

Supplemental Figure S2. Immunolocalization and quantification of DMC1 in zygotene meiocytes of Col-0 grown at 5, 12, 20 and 28°C.

Supplemental Figure S3. Dual-immunolocalization of SYN1 and DFO on prophase I chromosomes in Col-0 at 20°C.

Supplemental Figure S4. DAPI staining of pachytene, diakinesis and metaphase I chromosomes in the *dfo-1* mutant at 20°C.

Supplemental Figure S5. Dual-immunolocalization of SYN1 and DFO on prophase I chromosomes in *dfo-1* at 20°C.

Supplemental Figure S6. Expression analysis of the *DFO* gene in meiosis-staged flower buds in Col-0 at 20 and 37°C.

Supplemental Figure S7. Immunolocalization and quantification of ɤH2A.X in meiocytes of Col-0 and *spo11-1-3* at 20 and 37°C.

Supplemental Figure S8. Immunolocalization and quantification of ɤH2A.X on zygotene chromosomes of Col-0 and *atm-2* at 20°C.

Supplemental Figure S9. Immunolocalization and quantification of ɤH2A.X on zygotene chromosomes of Col-0 and *mre11-3* at 20°C.

Supplemental Figure S10. Meiotic and mitotic chromosome behaviors in meiocytes of *spo11-1-3 mre11-3* at 20 and 37°C.

Supplemental Figure S11. Quantification of mitosis defects in Col-0, *atm-2* and *mre11-3* at 20 and 37°C.

Supplemental Figure S12. Analysis of homolog synapsis and SC formation in *mre11-3* at 20 and 37°C.

Supplemental Table S1. Primers used in this study.

## Acknowledgment

The authors appreciate Dr. Wojtek Pawlowski (Cornell University) critical comments on the original manuscript prior to submission. They thank Dr. Shunping Yan (Huazhong Agricultural University) for kindly providing the seeds of the *atm-2, atr, asap1, sni1-2, asap1 rad51* and *sni1-2 rad51* mutants. They thank Dr. Yingxiang Wang (Fudan University) for kindly sharing the *mre11-3* mutant and the vector used for expressing the full-length AtRAD51 protein. They thank Dr. Zhongnan Yang (Shanghai Normal University) and Dr. Mathilde Grelon (Université Paris-Saclay) for sharing the seeds of the *dfo-1* mutant. They thank Dr. Holger Puchta (KIT) for sharing the seeds of the *spo11-1-3* mutant. They also thank GL Biochem, Shanghai, Ltd. (www.glschina.com) and Mabnus Ltd., Wuhan (www.mabnus.com/) for generating antibodies used in the study. In particular, Dr. Bing Liu appreciates Mrs. Meirong Huang, Mrs. Manxiang Zhu, Mr. Donglin Liu, Mrs. Jiajia Zhao and Ms. Zipei Liu for their support during the research.

